# Spatio-temporal patterns of wildfires in the Niassa Reserve –Mozambique, using remote sensing data

**DOI:** 10.1101/2020.01.16.908780

**Authors:** Eufrásio Nhongo, Denise Fontana, Laurindo Guasselli

## Abstract

Wildfires are among the biggest factors of ecosystem change. Knowledge of fire regime (fire frequency, severity, intensity, seasonality, and distribution pattern) is an important factor in wildfire management. This paper aims to analyze the spatiotemporal patterns of fires and burned areas in the Niassa Reserve between 2002-2015 using MODIS data, active fire product (MCD14ML) and burned area product (MCD64A1). For this, the annual and monthly frequencies, the trend of fires and the frequency by types of forest cover were statistically analyzed. For the analysis of the spatial dynamics of forest fires we used the Kernel density (Fixed Method). The results show a total of 20.449 forest fires and 171.067 km^2^ of burned areas in the period 2002-2015. Fire incidents were highest in 2015, while the largest burned areas were recorded in 2007. The relationship between increased fires and burned areas is not linear. There was a tendency for fires to increase, while for burnt areas there was stabilization. Forest fires start in May and end in December. August-October are the most frequent period, peaking in September. Fires occur predominantly in deciduous forests and mountain forests because of the type of vegetation and the amount of dry biomass. There is a monthly spatial dynamics of wildfires from east to west in the reserve. This behavior is dependent on vegetation cover type, fuel availability, and senescence.

## Introduction

Forest fires are among the biggest drivers of global ecosystem change. And one of the main forms of vegetation damage, threat to biodiversity, ecosystem weakening, soil infertility, production of harmful emission gases to human health, and decreased atmospheric visibility [1], [2].

In addition, biomass burning by fire has been identified as a significant source of aerosols, carbon fluxes and waste gases, which pollute the atmosphere and contribute to global climate change [3].

Despite these impacts, fire is a key ecological process in the miombo ecosystems, prevalent in the Niassa Reserve with impacts on nutrient cycling, vegetative regeneration, species composition, structure and resilience of the miombo ecosystem. According to [4], the fire in this phytophysiognomy is often labeled as a harmful phenomenon to ecosystems, but at the same time, it is an important management tool for ecosystems. Recent studies indicate increased recurrence of forest fires in the Niassa Reserve [5],[4], with serious economic impacts and the structure and composition of plant species. [6] show a reduction in the woody biomass of *Julbernardia globiflora* species in the Niassa Reserve between 2005 and 2009, and that this behavior may be linked to the increased frequency of burning.

Forest fire is a very dynamic landscape process that depends on different factors such as climate, vegetation type and structure, fuel moisture, land use and human activity [7].

Knowledge of the fire regime (fire frequency, severity, intensity, seasonality, and distribution pattern) is an important factor in forest fire management [8], [9]. Understanding and characterizing, for example, spatio-temporal fire-ignition patterns can provide important information for optimizing fire the allocation of resources [10],[11]. Therefore, in order to establish proper fire prevention policies, it is necessary to know the statistics related to them, i.e., to know where, when and why they occur [12].

In this context, the characterization of spatiotemporal patterns (fire regime) may contribute to improve the knowledge about how fires generally behave. Lack of this information can lead to significant expense, above the potential for damage, or very low spending [13].

Additionally, the description of the fire regime is an important factor in understanding the spatial distribution of plants and their regeneration strategies, particularly in regions with climatic seasonality [14].

In recent years, the availability of data from space satellites with different spatial, temporal and spectral resolutions has allowed access to raw data useful for detecting and monitoring active fires and extracting burnt debris not only at national or local level, but also at global scales [15]. These technologies allow comparison with other methodological approaches because of their accuracy, repeatability and speed of data acquisition, longer historical series and ease of combination with other thematic data [16].

Several sensors have been extensively used to study burned areas and fire outbreaks, including: Advanced Very High Resolution Radiometer (AVHRR); LANDSAT; Geostationary Operational Environmental Satellite (GOES); Medium Resolution Imaging Spectrometer (MERIS) and Moderate Resolution Imaging Spectroradiometer (MODIS) [17]. However, the timely resolution of MODIS, the availability of active fire and burned area products, the global scale and the fact that data is free of charge, has significantly increased the interest of global community end users in their adoption for regional and local applications, especially in areas where terrestrial data is scarce or not publicly accessible [18,19].

In this perspective, several studies have been developed to analyze the spatiotemporal patterns of forest fires based on global MODIS sensors [20], [21], [22], [23]. However, few studies on spatiotemporal patterns of fire occurrence have been conducted in the Niassa Reserve.

Thus, the present work aims to analyze the spatiotemporal patterns of forest fires, detected from the MODIS satellite, between 2002 and 2015, in the NR. The main question to be answered is where and when fires occur. Contributing in this way to the study of fires in the Niassa Reserve, as it is the first study that analyzes the spatial-monthly dynamics of forest fires in the Niassa Reserve.

## Materials and Methods

### Study Area

The study area is the Niassa Reserve (23.040 km^2^), located in the far north of Mozambique, between latitudes: 12 ° 36’46.67” - 11 ° 26’05.83” south, and longitude: 32 25’20.16” and 38 ° 31’23.16” east (Fig. 1). The reserve is part of the Rovuma watershed. The relief ranges between 136 and 1.413 m, above mean sea level. It is one of the largest conservation areas in southern Africa and the largest in Mozambique, and is one of the most iconic wilderness areas in Africa. [24]

**Figure 1:**
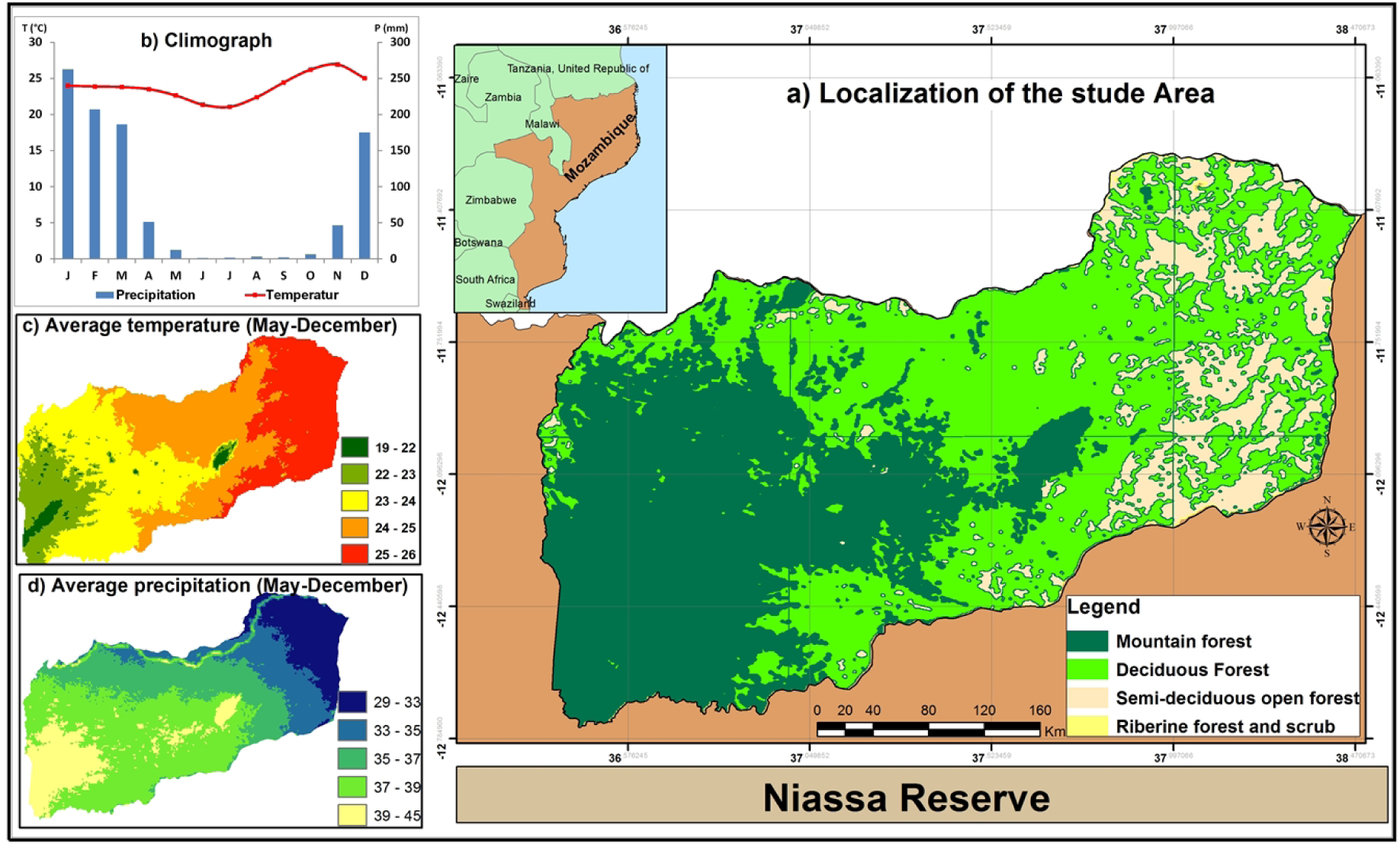
Geagraphic Location of Niassa Reserve: a) Homogeneous regions of vegetation cover according to NDVI temporal variability (Source: Nhongo et al., 2017); b) Climograph; c) Average Temperature (May-December); d) Average precipitation (May-December).

**Figure 2:**
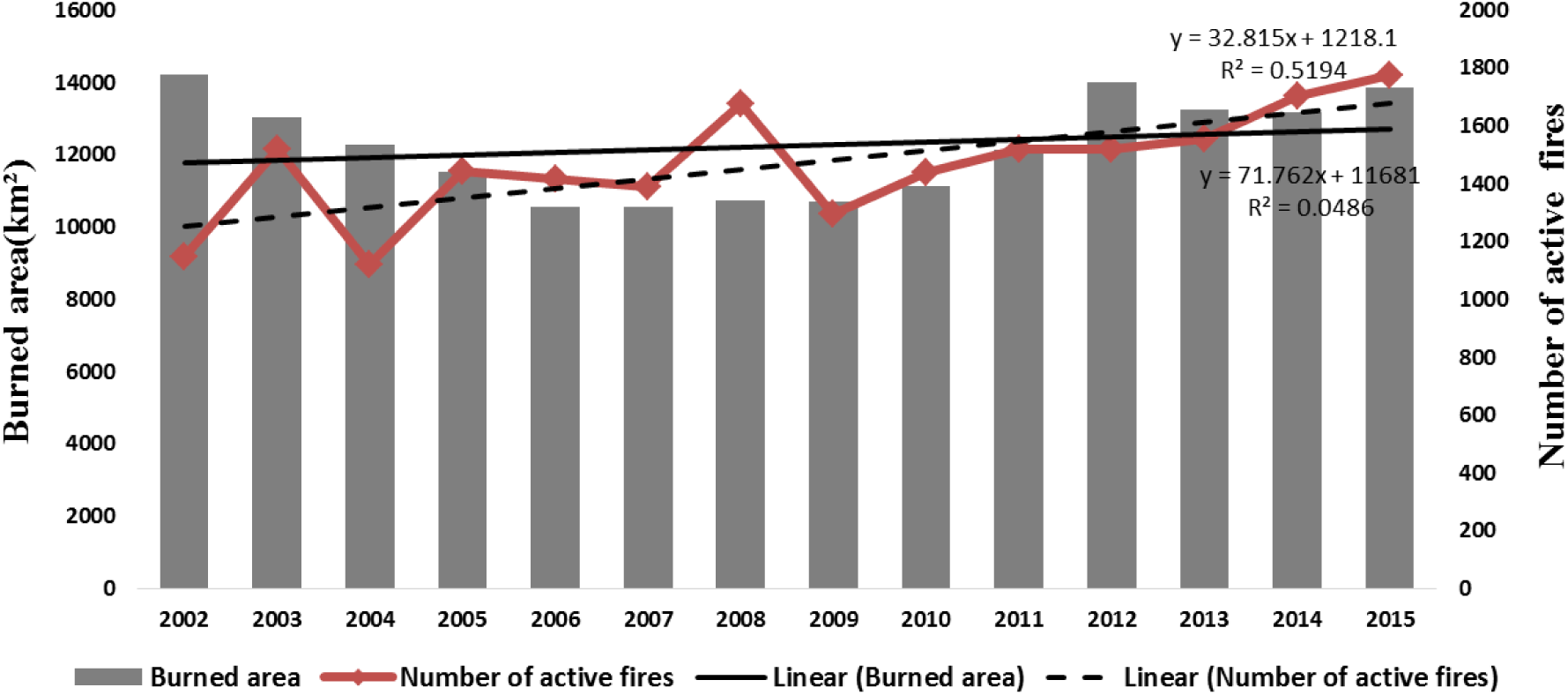
Burned area and number of wildfires (2002–2015), in Niassa National Reserve, The line represents linear trend: solid line for burned area, and dashed for number of fires.

The climate type is tropical sub-humid, with two distinct climatic seasons: dry (May to September) with rainfall not exceeding 20 mm, and relative humidity between 40 to 50%, this is the period with the highest occurrence of forest fires.

According to [25], the predominant vegetal formations are: Deciduous forest, Semi-deciduous open forest, Mountain forest, and Riverine forest and Scrub.

## Data

### MODIS MCD14ML Active Fire Product

Were obtained from the Aqua & Terra MODIS thermal anomaly, with spatial resolution of 1 km, Collection 6, from January 2002 to December 2015, available from NASA’s Fire Information for Resource Management System (FIRMS) website services (https://earthdata.nasa.gov/earth-observation-data/near-realtime/firms/)

Fire detection is performed using a contextual algorithm that exploits the strong emission of medium infrared fire radiation and is based on the brightness temperature derived from the 4 and 11 µm channels. [26]. The location of the fire corresponds to the center of a 1×1 km pixel, signaled by the algorithm as containing one or more fires within the pixel. To avoid false alarms (commission errors), only highly reliable fire pixels (> 80% reliability) were considered.

### Product of burned area MODIS MCD64A1

The burned area data were obtained from the MODIS burned area sensor, monthly product MCD64A1 Version 6, available from the MODIS Active fire and Burned Area products website (http://modis-fire.umd.edu/). MCD64 is the latest product from the MODIS Burned Area product. This is a 500 m product, global grid level 3. It is based on an automated hybrid approach that exploits the 1 km MODIS active fire potential and 500 m surface reflectance input data [27].

The hybrid algorithm applies dynamic boundaries to composite images generated from a burn sensitive vegetation index, which in turn are derived from shortwave infrared channels, MODIS band 5 and 7, and a measure of temporal texture. Data layers include recording date, recording data uncertainty, quality assurance and the first and last day of reliable change detection of the year

Overall the MCD64A1 has improved detection of burned areas over past collections. But specifically in significantly better detection of small fires and adaptability to different regional conditions in multiple ecosystems. [28].

## Method

### Temporal Pattern

In order to obtain the spatial-temporal distribution pattern of Niassa Reserve fires, the time series of fires and burned areas (2002-2015) was analyzed using ArcMap 10.1 and descriptive statistics. Statistical analysis advocated: (1) Annual frequency analysis; (2) Linear trend analysis; and (3) Monthly frequency analysis.

### Spatio-temporal distribution pattern

#### Distribution by vegetation cover

The frequency analysis was complemented by crossing fire outbreaks with different types of forest cover in the reserve, seeking to identify the forest cover with the highest occurrence of fire, total burned area and the monthly dynamics.

The forest cover map used is the map Homogeneous regions of vegetation cover according to NDVI temporal variability prepared by [25]. This is important to identify the vegetation cover patterns most affected by forest fires. It should be noted that forest cover types where forest fires and burned areas were not recorded were not used, such as the riverine forest and scrub.

#### Kernel Density Estimation

For analysis of the monthly spatial pattern of fires, kernel density was applied. Kernel estimator is a nonparametric statistical method that produces a smoothed cumulative density function. [29, 30] effectively used to map fire occurrences [31]. Kernel estimator is mathematically defined [32, 33] by the equation 4:

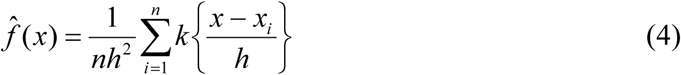

Where: *n* is the number of points observed; *h* is the bandwidth; *K* is the Kernel function; *x* is the coordinate vector representing the location of the estimated point; and *Xi* is the vector of the ith coordinate that represents each point observed in relation to the estimated.

One of the key steps in estimating kernel density, in addition to choosing the kernel function, is setting parameter smoothing, such as bandwidth size (‘search radius’ in ArcGIS 10.2). Often determined on the basis of subjective choices, specialized knowledge or possibly supported by empirical decisions.

According to [32], the choice of bandwidth depends on the purpose of density estimation. If the goal is to explore the data and suggest models and hypotheses about it, it is sufficient to choose the smoothing parameter subjectively by visual inspection. However, it is very difficult to define this subjective value, which can generate ambiguous results, because the values depend on the scale adopted and the specific characteristics of the studied area [31], [34]. There are two main methods used to find the appropriate bandwidth size: Fixed and Adaptive. The fixed method is used in regions where data is dense and evenly distributed, without evidence of clustering [34] and distance units, is constant throughout the area of interest. In adaptive mode, the smoothing parameter, defined using a minimum number of point observations found under the nucleus, varies depending on the concentration of point observations [35].

Since there is spatial variability of data in the reserve, and the total fire distribution is regular or homogeneous with respect to the total reserve area, both methods could be adopted. However, for reasons of comparing the monthly density, we opted for fixed bandwidth. For this we used the mean distance method (RDmean), which can be analyzed by local or global approach. Defined by [31] as:

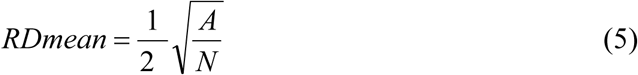

Where: <u>A</u> is the total size of the study area; <u>N</u> is the total number of Heat Focuses.

The parameters used and the bandwidth result are presented in Table 1.

**Table 1:** Parameters Related to Bandwidth Calculations.

|  |  |
| --- | --- |
| Total area of study, A | 22.953km<sup>2</sup> |
| Total number of polygons | 3 |
| Total number of active fires, N | 20.499 |
| Polygon Size Average | 7.651 Km<sup>2</sup> |
| Average number of fires active per polygon | 7.074 Km<sup>2</sup> |
| RDmean Local 2x | 1.059,4 m |
| RDmean Global 2x | 1.059,4 m |

## Results

### Temporal Patterns

From the MODIS images, a total of 20.449 forest fires were recorded between 2002 and 2015, an annual average of 1,464 fire outbreaks and 171,067 km^2^ of burned areas, an annual average of 12.219,1 km^2^ of area, equivalent to 53.2% of the total area of the Niassa Reserve.

The year with the highest record of forest fire outbreaks was 2015 with 1774 fires detected, and the year of greatest severity (with the largest burned area) was 2002, with about 14.200 km^2^. Overall, there was a tendency for the number of forest fires increase in the series analysis, while burnt areas show no upward or downward trend over 14 years.

The total monthly distribution chart of forest fires and burned areas comprises the historical series with accumulated monthly totals from 2002 to 2015. (Fig. 3).

**Figure 3:**
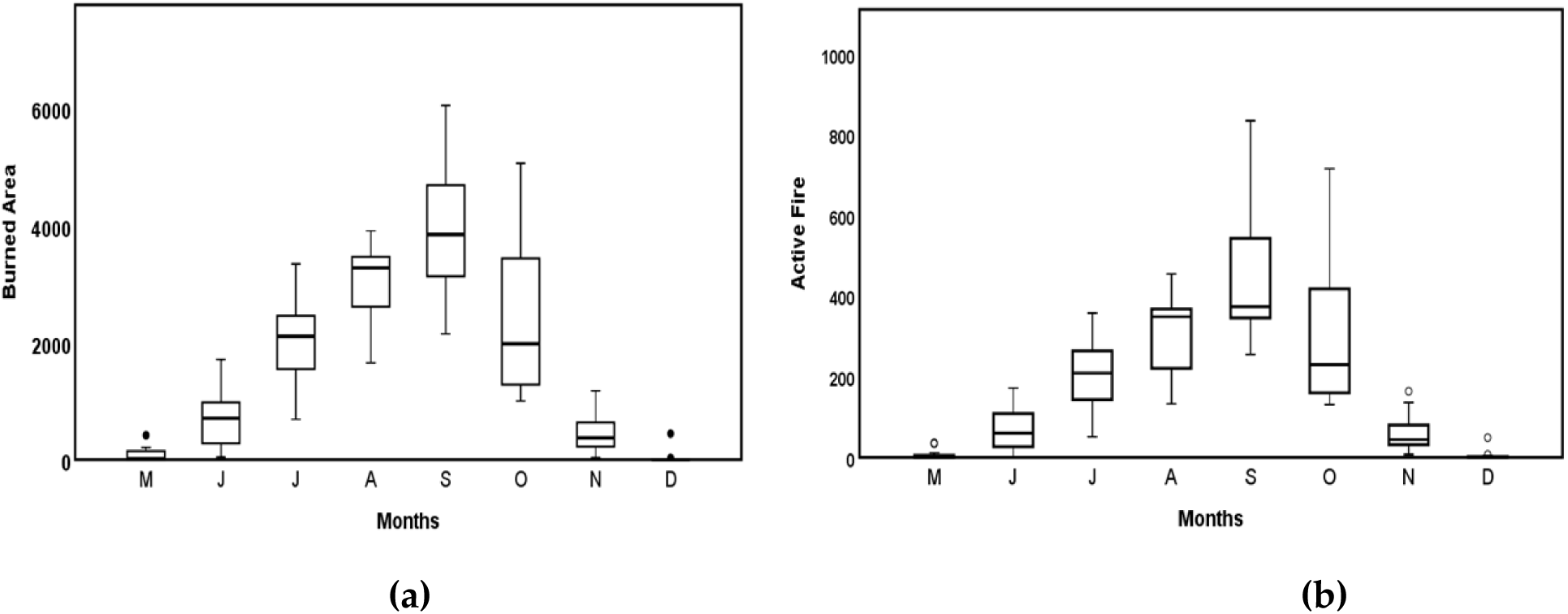
Monthly variation in wildfire occurrence in Niassa Reserve: (a) Burned Area(km^2^).; and (b) Active Fire

The occurrence of fires and burned areas was recorded between May and December, the months with the highest occurrence of forest fires are August, September and October. However, September is the month with the highest record of fires and burned areas, with 32.6% of fire outbreaks and 31.60% of burnt areas.

### Spatial patterns

The spatial pattern of fires was influenced by forest cover types. The fires occurred predominantly in areas of deciduous forest (50%) and mountain forest (38%). Burned areas followed the same trend in deciduous vegetation (51%) and Mountain forest (35%). Semi-deciduous open forests have fewer occurrences of forest fires (Tabela 2).

**Table 2:**
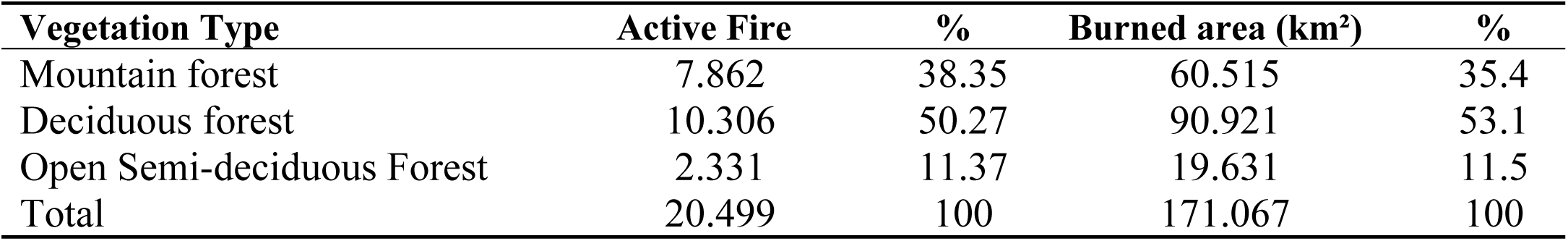
Total distribution of number of fires and burned areas by forest cover 2002-2015, Niassa Reserve.

| Vegetation Type | Active Fire | % | Burned area (km<sup>2</sup>) | % |
| --- | --- | --- | --- | --- |
| Mountain forest | 7.862 | 38.35 | 60.515 | 35.4 |
| Deciduous forest | 10.306 | 50.27 | 90.921 | 53.1 |
| Open Semi-deciduous Forest | 2.331 | 11.37 | 19.631 | 11.5 |
| Total | 20.499 | 100 | 171.067 | 100 |

It is noteworthy that there is a spatial-monthly dynamics of outbreaks and burnt areas in the reserve. From May to June (early burning season), fires start east of the reserve (Fig. 5), with the highest occurrence of fires and burned areas in semi-deciduous open forests (Fig 4), and migrate gradually to the center of the Reserve.

**Figura 4:**
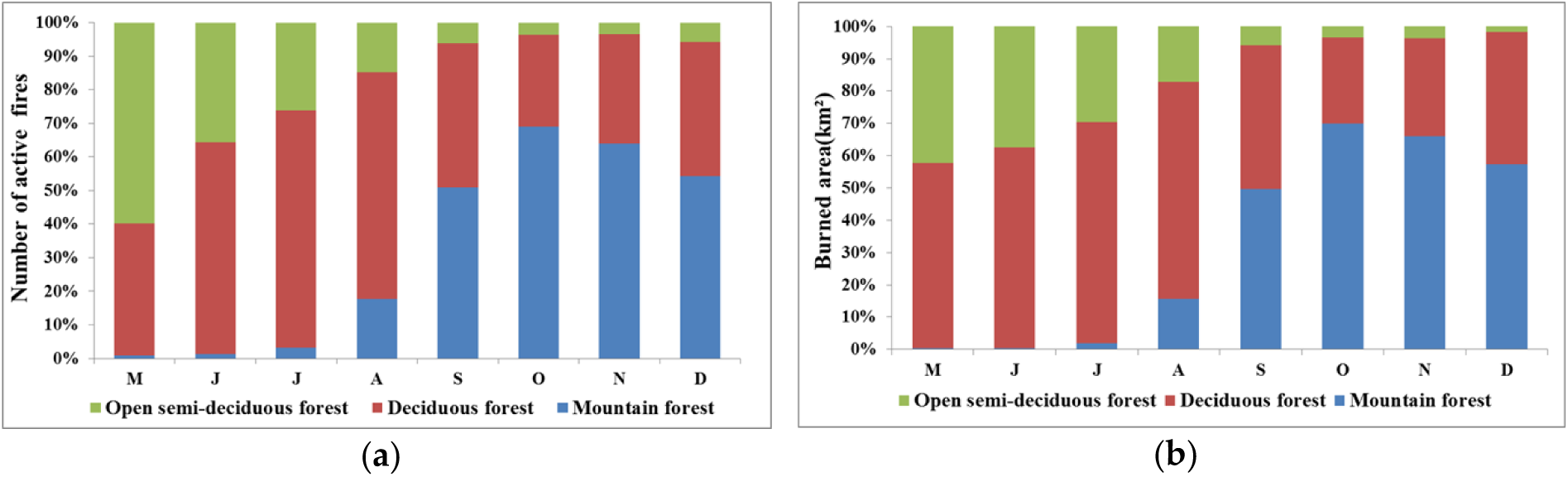
**a)** Monthly distribution of number of fires by vegetation cover; b) Monthly distribution of burned areas by vegetation cover.

**Figure 5:**
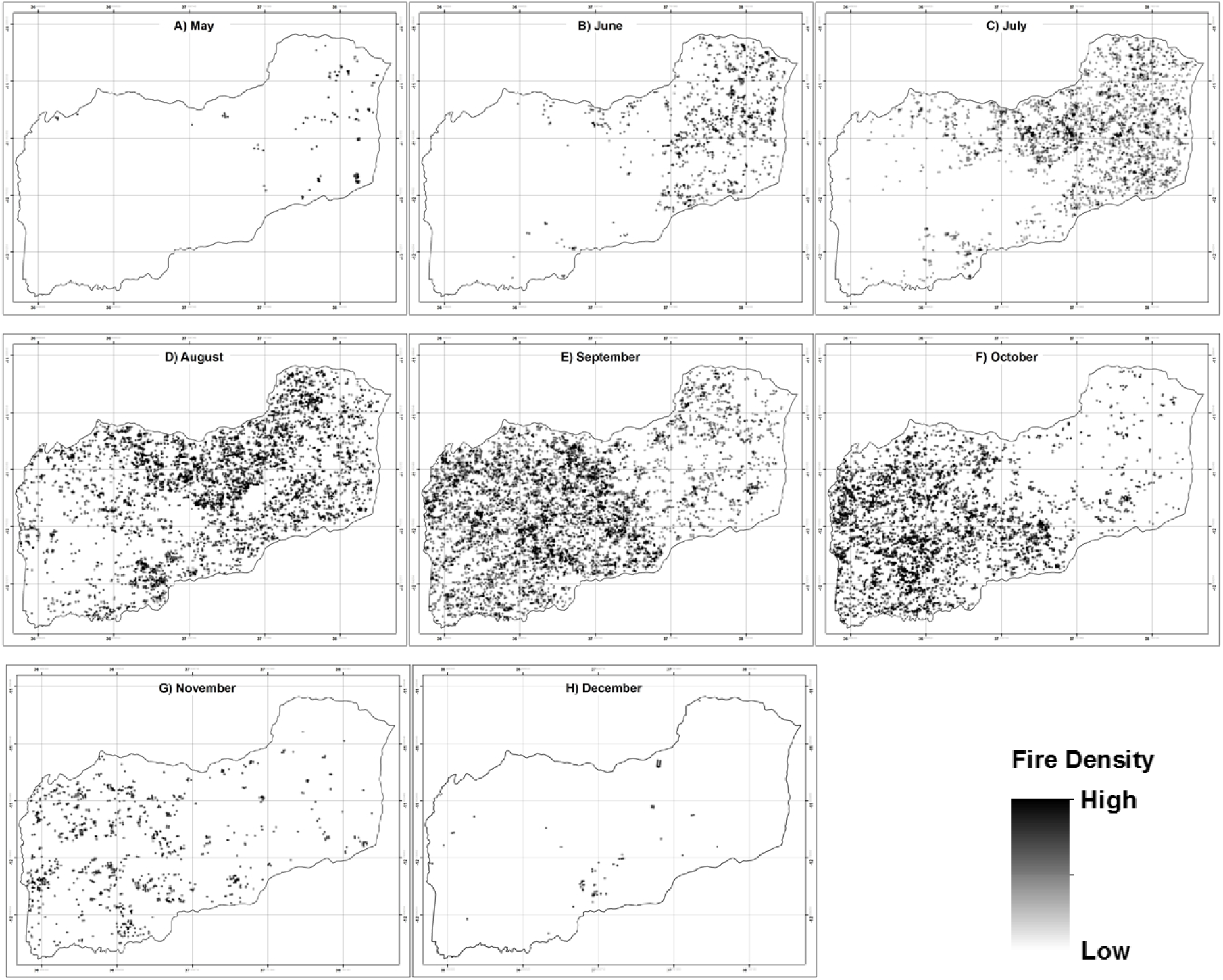
Spatial patterns of wildfire density from May to December, Niassa Reserve, Mozambique.

From July to August there is a migration and predominance of fires and burned areas in deciduous forests (Fig. 4A and 4B). Especially in the central east, central and midwest regions (Fig. 5). However, in September, fires migrate and predominate in deciduous forests as well as mountain forests (Figure 4A and 4B), and are located in the center and west of the reserve (Figure 5). From October to December there is a migration of fires to the west of the Niassa Reserve, with a predominance of Mountain forests.

## Discussion

### Temporal Pattern

This study provides evidence that the occurrence of fires in the Niassa Reserve are temporally and spatially variable. The highest peak of fire occurred in 2015 and the highest severity in 2002.

However, the results show an increase in fires that resulted in large areas burned between 2012-2015, which may be related to meteorological phenomena such as temperature increase and low rainfall over the analyzed period

The occurrence of fires and burned areas was recorded between May and December, with the peak recorded in September (Fig 3). These results are consistent with studies conducted in southern Africa [36], [37], [38].

The burning period begins in May, when the deciduous open vegetation of the reserve goes into senescence [25]. This is not surprising, as most fires occur during the dry season, after the senescence period in various phytophysiognomies around the world [38], [39], [40].

It is also observed that the highest occurrence of fire and the highest severity occur between August, September and October, with about 70% of fire and 73.6% of burned area, with maximum peak in September. (Fig 3). What can be explained by a combination of factors, September is the month with the lowest rainfall (Fig. 1b), low relative humidity, and highest dry biomass accumulation, which causes greater ignition occurrence and ease of propagation fire, especially in deciduous and mountain vegetation. Similar results were found in studies conducted on similar phytophysiognomies in southern Africa [38], [41], [42].

The end of the fires occurs between November and December, when rainfall falls on average exceed 150 mm (Fig. 3). This result was not expected, as the hydrological year begins in October. However, according to [13] the increase in precipitation does not necessarily correspond to the immediate reduction in the occurrence of fires. As the soil and combustible material are still low in moisture, precipitation will be absorbed to the point where the combustible material will no longer ignite. However, October is the month with the highest amplitude or variability of fire occurrence in the reserve, which may be related to the beginning of the hydrological year in the second half of this month.

### Spatial pattern

The spatial pattern of fires was influenced by the type of forest cover. The fires occurred predominantly in areas of deciduous vegetation and mountain forest. The burned areas followed the same trend with greater severity in deciduous vegetation and mountain forest. (Fig. 2).

These results were expected, as approximately 60% of the reserve is occupied by deciduous forests, and the phytophysiognomy presents very low values of vegetation vigor in the dry period. However, Mountain forests were not expected to have high numbers of forest fires and burned areas, which may be related to the greater accumulation of dry biomass, especially from September to October.

It is noteworthy that there is a spatial-monthly dynamics of outbreaks and burnt areas in the reserve. From May to June (early burning season), fires start east of the reserve, with the highest occurrence of fires and burned areas located in semi-deciduous open forests (Figs. 4 and 5).

The main factors that determine the beginning of the fire season in this region and in this phytophysiognomy are the high temperature and the low precipitation. According to [25] this is the first phytophysiognomy that enters the senescence process in the Niassa Reserve and consequently higher probability of fire in this period.

According to [43] fires are limited by the availability of fine fuel, especially in semi-deciduous open forests, which in turn are dependent on soil moisture and nutrient availability. Thus, the low amount of dry biomass in Semi-deciduous Open Forests (Eastern Reserve Region) determines the gradual (spatial-temporal) migration of fires to the Reserve Center in deciduous forests (Fig 5).

From June to August there is a migration and predominance of fires and burned areas in the deciduous forests (Figs 4 and 5), especially in the central east, central and midwest regions. In this phytophysiognomy, in this period, there is the beginning of senescence, and consequent reduction of vegetation vigor [25].

With the end of dry biomass in deciduous forests and the onset of senescence in mountain forests, starting in September, the same process of migration of fires to mountain forests west of the reserve occurs.

This region covered by wetter forests, with a long phenological cycle, requires more time to provide flammable conditions for forest fires. The trend of increased occurrence of fire outbreaks and burned areas in this phytophysiognomy continues until December.

According to [44], fires migrate when there is no longer enough fuel to sustain them, when weather conditions are not prone to burning or when they encounter topographic or anthropogenic barriers or previously burned areas. It is evident in the reserve area that the lack of sufficient fuel for its sustainability is fundamental to the dynamics of fires. But, as already mentioned, the beginning of the burnings and their migration are also strongly dependent on senescence and the availability of dry fuel.

### Conclusions

This study aimed at analyzing the spatiotemporal patterns of forest fires in the Niassa Reserve based on MODIS active fire product (MCD14ML) and burned area product (MCD64A1) data.

- The existence of a spatial-temporal, monthly and annual pattern of fires and burned areas in the Niassa Reserve was observed.
- There is a tendency for fires to increase, but this increase has not resulted in increased burned areas.
- The burning season begins in May and ends in December. The period of greatest fire occurrence is from August to October, with a peak in September.
- Deciduous forests and mountain forests have the highest occurrence of fires, due to their structure and dry biomass accumulation.
- There is a monthly spatial dynamics of wildfires from east to west of the reserve, which is strongly dependent on the type of vegetation cover and dry biomass accumulation.
- Fire data from MODIS products can be used successfully to understand the spatial and temporal extent and distribution of fire activities and in different vegetation types.

## Author Contributions

EJSN. Provided the ideal and constructive suggestions towards the whole project performed the data processing as well as wrote the manuscript. DCF Provided the ideal and constructive suggestions towards the whole project, as well as wrote the manuscript. LG Provided the ideal and constructive suggestions towards the whole project, as well as wrote the manuscript.

## Acknowledgments

To the National Council for Scientific and Technological Development (CNPq), for granting a PhD scholarship to the first author. The Ministry of Science and Technology of Mozambique for funding the publication of the Scientific article.

